# Natural disturbances and connectivity shape the seasonal variability of aquatic macroinvertebrate communities across Europe

**DOI:** 10.1101/2024.10.09.617360

**Authors:** Loic Chalmandrier, David Cunillera-Montcusi, Naiara Lopez-Rojo, Miguel Canedo-Arguelles, Zoltan Csabai, Arnaud Foulquier, Franck Jabot, Marko Milisa, Heikki Mykra, Petr Paril, Balint Pernecker, Luka Polovic, Romain Sarremejane, Henna Snare, Maria Soria, Thibault Datry, Nuria Bonada, Francois Munoz

**Affiliations:** University Grenoble-Alpes; Center for Ecological Research; INRAE Riverly; Institute of Environmental Assessment and Water Research; University of Pecs, Department of Hydrobiology; INRAE; University of Zagreb; Finnish Environment Institute; Masaryk University; University of Pecs; Nottingham Trent University; Universitat de Barcelona; University of Barcelona; University Grenoble Alpes

**Keywords:** Communities, River drying, Connectivity, Seasonal dynamic, Traits, Macroinvertebrate, Dispersal, Disturbance, Resistance, Resilience

## Abstract

Understanding the joint influence of natural disturbance, spatial connectivity and biogeography on biodiversity is essential to forecast its responses to climate change. Macroinvertebrate communities in drying river networks constitute an ideal study system to understand the interplay of these ecological processes. We analyze the taxonomic and functional structure of macroinvertebrate communities sampled across 126 reaches with perennial and intermittent streamflow, surveyed in six drying river networks (DRN) across Europe, six times over one year.

Drying frequency decreased community richness and functional diversity of communities, whereas spatio-temporal connectivity increased community richness in intermittent reaches. Communities experiencing a high drying frequency increased the proportion of taxa with K-strategies and drying resistance traits. Communities experiencing a long drying duration compensated by high spatio-temporal connectivity had more taxa with a r-strategy and high dispersal ability. Perennial communities varied from taxa-poor communities of r-strategists in spring and autumn and taxa-rich communities of K-strategists in summer and had a constant functional diversity throughout the year. When drying frequency increased, communities showed a similar pattern except in autumn when they shifted towards species-poor communities of K-strategists. Functional diversity then peaked in summer. Community trait structure and in particular optimal drying resistance traits changed across biogeographical scales. It opposed communities from mountainous DRN (with more r-strategies and high dispersal ability) to non-mountainous DRN (with more K-strategies).

Drying frequency, drying duration, and spatio-temporal connectivity drive divergent community structures, suggesting the presence of an ecological threshold that explains the variability of disturbed ecosystems across broad spatial scales. These factors also shaped seasonal community variations, particularly after summer, with intermittent communities influenced by stochastic recolonization events in spring and autumn. Spatial-temporal connectivity proved crucial for maintaining diversity in communities subjected to intense drying. Lastly, the effectiveness of drying resistance traits was dependent on the biogeographical and environmental conditions of drying river networks.

## Introduction

Global change is profoundly altering natural disturbance regimes and the seasonal variability in climate, with dramatic implications for biological communities (Harris et al., 2018; Heino et al., 2009; Pau et al., 2011). Natural disturbances, defined here as recurring events that cause a sudden increase in population mortality (Jentsch & White, 2019; Miller et al., 2011) alter niche processes by modifying the abiotic stresses experienced by species and further alter dispersal processes by disrupting habitat connectivity (Jacquet et al., 2022; Tonkin et al., 2018). Seasonal variability causes temporal fluctuations in environmental conditions, affecting species’ life cycles and interactions (Mellard et al., 2019; Pau et al., 2011). Integrating these factors in the analysis of communities is essential to accurately forecast biodiversity responses to climate change (Holyoak et al., 2020; Tonkin et al., 2017).

Drying river networks (DRNs) provide an ideal system for studying the complex interplay of these ecological processes. Representing a significant portion of global river systems, DRNs play a crucial role in maintaining biodiversity and ecosystem functions, yet are increasingly threatened by global changes (Datry et al., 2023; Messager et al., 2021). These networks contain intermittent reaches, where flow cessation and resumption are shaped by natural disturbances, i.e. drying events. The frequency and duration of these drying events create a dynamic mosaic of wet and dry patches, which can fragment the network and reduce hydrological connectivity between wet patches (Cañedo□Argüelles et al., 2015; Cunillera-Montcusí et al., 2023a). As a result, DRN communities are influenced by both stochastic and deterministic processes. Intermittent reaches, compared to perennial ones, tend to host species-poor, randomly assembled communities due to local extinctions during drying events and stochastic recolonization upon flow resumption (Sarremejane, Mykrä, et al., 2017). These effects are exacerbated when the loss of hydrological connectivity limits dispersal within the network, promoting ecological drift (Carrara et al., 2014; Jacquet et al., 2022).

Differences in drying and connectivity regimes across river reaches are expected to influence the functional structure of macroinvertebrate communities in a deterministic manner (Crabot, Mondy, et al., 2021; Crabot, Polášek, et al., 2021; Gauthier et al., 2020). Aquatic macroinvertebrates, for instance, may endure dry phases through diapause or resistant life stages (e.g., eggs, cocoons). This selection of species with adequate resistance/resilience traits should decrease functional diversity in intermittent communities compared to perennial ones. Conversely, species may exhibit resilience traits (e.g. high dispersal ability, high fecundity or short generation times) that allow for rapid recolonization once flow resumes if the community is well connected to the rest of the river network (Datry et al., 2016; Jabot et al., 2020; Sarremejane, Cañedo□Argüelles, et al., 2017). Thus, intermittent reaches with greater connectivity to the river network should have a higher proportion of species with resilience traits coexisting with species with resistance traits and thus functional diversity should increase (Cunillera-Montcusí et al., 2023b).

Significant knowledge gaps remain regarding how DRN community assembly varies across different drying regimes and biogeographical scales (but see Crabot, Mondy, et al., 2021; Vander Vorste et al., 2021). First, variations in community assembly may result from differences in intermittence regimes, as intermittent reaches can vary in both the frequency and duration of drying events. These variations are often driven by climatic factors—such as the prolonged and frequent drying events characteristic of Mediterranean climates (Bonada et al., 2007; Tonkin et al., 2017) —or by geomorphological differences (Foulquier et al., 2024; Snelder et al., 2013; Vander Vorste et al., 2021). However, few studies have systematically examined the distinct effects of drying frequency versus duration on the taxonomic and functional structure of macroinvertebrate metacommunities, or how these factors contribute to biodiversity turnover across large spatial scales. Second, differences in community structure across DRNs may also be influenced by varying evolutionary legacies (Bonada et al., 2007, Bonada et al., 2017). For example, Mediterranean DRNs likely support a higher proportion of taxa with resilience or resistance strategies, as local species have evolved under recurrent intermittent regimes since at least the last glaciation.

Seasonal changes in river discharge, water temperature, and oxygen levels cause seasonal shifts in community structure of river communities (Tonkin et al., 2017). Research gaps persist regarding the seasonal dynamics of freshwater macroinvertebrate communities, particularly in relation to shifts in trait structure (Woods et al., 2022 but see Novella-Fernandez et al., 2023). Broad ecological expectations suggest that taxonomic diversity peaks in summer when cold-related stress is minimal and that communities are shaped more by abiotic filtering in spring and biotic interactions in summer, leading to higher functional diversity in summer (Mellard et al., 2019; Pau et al., 2011). However, drying river networks may deviate from these patterns. Even if an intermittent reach does not fully dry, summer conditions can lead to critically low or stagnant discharge, creating abiotic stress due to elevated temperatures and low oxygen levels (Bonada et al., 2020). Consequently, communities in intermittent reaches may display lower-than-expected functional diversity during the summer (drying season), as higher abiotic stress selects for similar resistance traits (Stubbington et al., 2017). Additionally, many species adjust to the seasonal hydrological variation through short and multivoltine life cycles that commonly lead to peak insect emergence in early summer, before intermittent reaches are dry (Strachan et al., 2015).

While substantial research exists on macroinvertebrate communities in DRNs, current studies often have limited spatial and temporal scope and rarely address the combined effects of intermittence, river connectivity, and seasonality across different biogeographical contexts. To address these gaps and establish general community assembly rules in DRNs, we sampled macroinvertebrate communities six times over one year across 126 reaches in six DRNs across Europe. This extensive coverage of intermittence and seasonal variation allowed us to investigate how various aspects of river drying regimes—such as drying frequency, average drying duration, and spatio-temporal connectivity—influence the seasonal dynamics of macroinvertebrate functional and taxonomic community structures across Europe. This research will help identify both general and context-specific mechanisms driving community assembly in DRNs. Specifically, we test four hypotheses:

1. Communities in intermittent reaches will have lower individual density, reduced taxonomic and a distinct functional structure compared to perennial reaches, reflecting a higher prevalence of organisms with resistance and/or resilience strategies.
2. Intermittent reaches with greater connectivity to the broader river network will exhibit higher abundance, richness, and functional diversity, and a greater proportion of species with resilience strategies linked to strong dispersal capabilities than intermittent reaches with a low connectivity.
3. Seasonal variation in macroinvertebrate communities will be more pronounced in intermittent reaches compared to perennial ones. In summer, perennial reaches are expected to have higher functional diversity due to stronger biotic filtering, while intermittent reaches may show lower functional diversity due to the stress from low flow and high drying probability, selecting for species with resistance traits.
4. Community structure will differ between DRNs in Mediterranean and temperate climates. DRNs in Mediterranean regions are expected to host a higher proportion of taxa with resilience and resistance strategies compared to temperate DRNs.

## Methods

### Data

#### Sites and sampling strategy

This study is part of the project DRYvER [Securing biodiversity, functional integrity, and ecosystem services in DRYing riVER networks [DRYvER (Datry et al. 2021)]. The research was conducted in six DRNs across Europe, spanning longitudes from −5.328 to 24.656 and latitudes from 36.430 to 60.481. Sites encompassed five climatic zones (Figure 1): Mediterranean (Spain and Croatia), Continental humid (Czechia), Cold temperate climate (Finland), Pre-alpine climate (France) and Continental climate (Hungary). The study also covered six biogeographical ecoregions: Balkanic (Croatia), Continental (Czechia), Boreal (Finland), Continental/Alpine (France), HU Pannonian (Hungary), and Mediterranean (Spain). In each DRN, 15 to 26 reaches were selected to represent the whole river network, including both the main stream and tributaries, with a balanced selection of perennial and intermittent reaches. This classification was based on expert knowledge from the scientific teams surveying each DRN. Hydrological modeling later revealed that some surveyed perennial reaches could undergo drying events but at a much lower frequency and shorter duration than intermittent reaches (Figure S4).

**Figure 1.**
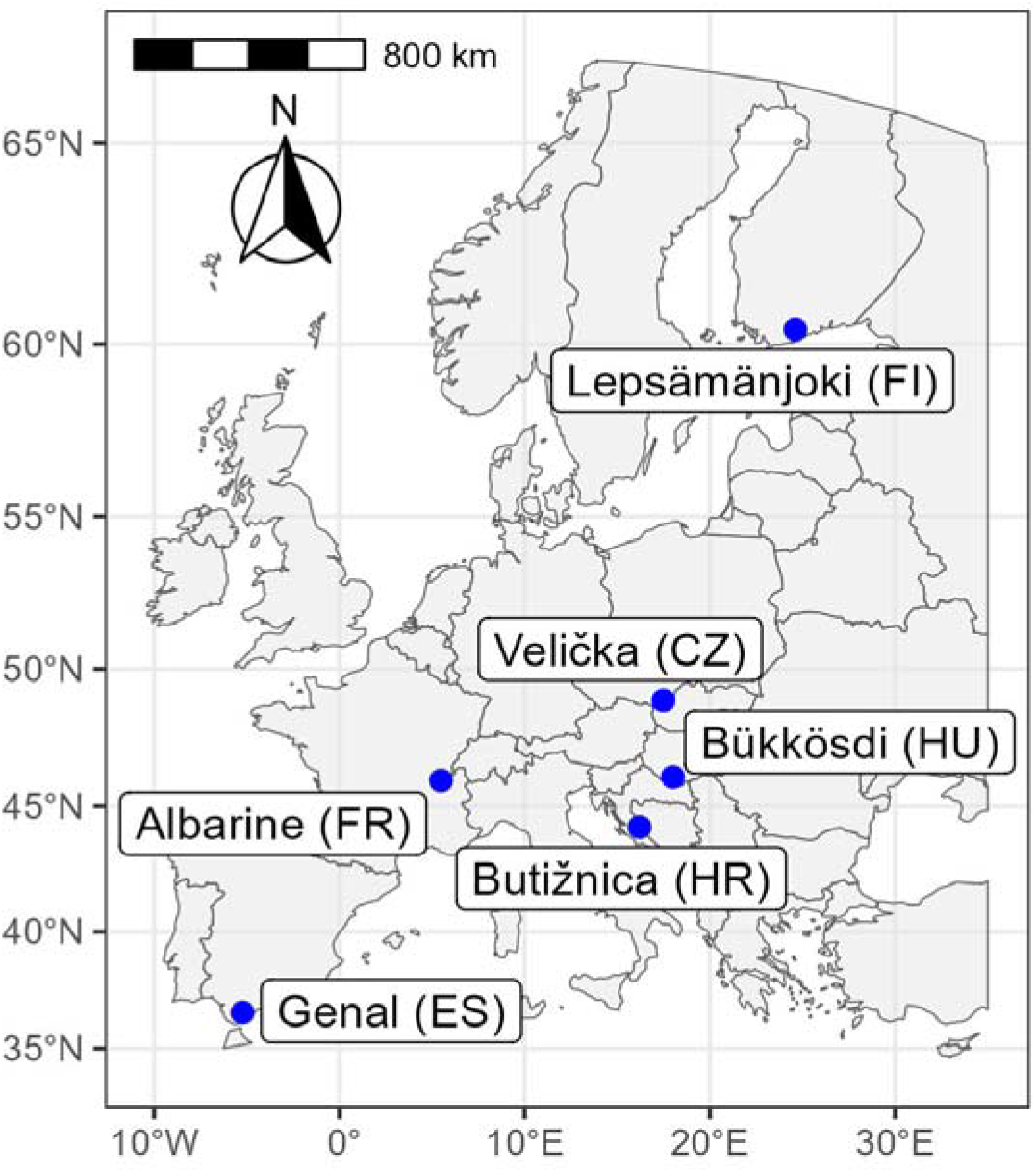
Location of the six drying river networks in Croatia (HR), Czechia (CZ), Finland (FI), France (FR), Hungary (HU) and Spain (SP).

Reaches with obvious anthropogenic hydromorphological alterations (e.g., channelized reaches, reaches located upstream/downstream of dams) were excluded to focus on the effect of natural drying events. Each sampling reach consisted of a river segment of at least 50 m and 20 times the maximum wetted width (measured in the first sampling campaign and held constant for subsequent ones) to cover several representative sequences of geomorphic units (e.g., riffles, pools) (Datry et al., 2015). See Figure S1 and S2 for an analysis of the environmental variation among and within sampling reaches.

#### Hydrological data

In each DRN, flow intermittence was modeled at the scale of the whole catchment by using a hybrid modeling approach (see details in Mimeau et al., 2024). Using this modeling work, we estimated three descriptors of intermittence for each community:

- Drying duration was calculated across all drying events in the 10 years preceding community sampling
- Drying frequency calculated as the number of drying events in the 10 years preceding community sampling
- Spatio-temporal river connectivity in the 30 days preceding each sampling campaign was estimated using the framework proposed in Cunillera-Montcusí et al., (2023).

The distributions of drying frequency and drying duration were right-skewed, so we apply a square-root transformation to their values for all analyses. In the supplementary materials, we assessed the covariation of spatio-temporal connectivity, drying duration and drying frequency across the year drying river network and reach type (intermittent vs. perennial).

#### Community data

Macroinvertebrates were sampled every two months in each DRN between January 2021 and February 2022 for a total of six sampled dates per DRN unless the reach was dry. The samples were identified as often as possible to genus level and we excluded terrestrial taxa from the analysis. Details about the sampling and identification process are available in the Supplementary materials.

#### Trait data

We grouped taxa into four groups representing contrasted biological traits, as often used in aquatic ecology: EPT (Ephemeroptera, Plecoptera, Trichoptera - 112 taxa), OCH (Odonata, Coleoptera, Hemiptera - 75 taxa), OI (Other insects, here grouping mainly Diptera and Megaloptera - 31 taxa) and finally NI (Non-Insects, including among others Mollusca, Crustacea and Annelida - 72 taxa). We constructed a trait database from two overlapping genus-level databases (Sarremejane et al., 2020; Tachet et al., 2010). We selected life-history traits and known resistance and resilience traits based on the overview provided by Stubbington et al., (2017): five were continuous (maximum body size, longevity, lifelong fecundity, drift, number of reproductive cycles per year) and two were categorical (dispersal mode - passive aquatic dispersal, active aquatic dispersal, passive aerial dispersal, active aquatic dispersal; and resistance forms - eggs; cocoons; diapause/dormancy; none).

In total, across 638 community samples, we sorted and identified over 1.35 millions aquatic macroinvertebrate individuals belonging to 290 taxa (240 at the genus level, 45 at the family and 6 at a higher taxonomic level). For trait-based analyses, we analyzed 368 community samples containing a total of 1.3 × 10^6^ individuals classified in 265 taxons with sufficient trait data (see Supplementary materials for details about the trait database construction).

### Data analysis

#### Community statistics

We analyzed total community density (as total number of individuals) and community taxa richness, community average trait structure and functional diversity for each community. For abundance-based metrics (except total abundance), individual counts in the site by taxa matrix were transformed with the function log(x+1). This was justified by the very high local counts of individuals for some taxa (e.g. *Gammarus* individuals could reach up to 30 000 individuals per sample) that would have a disproportionate influence on abundance-weighted community metrics.

We adopted a multivariate approach to study how the structure of communities varied in a multi-trait space. We first computed a correspondence analysis (CA) on the matrix of taxa relative abundances. We then computed a principal component analysis (PCA) on the trait matrix. While maximum body size, longevity, lifelong fecundity, drift, number of reproductive cycles per year were continuous traits (see above), dispersal and resistance were still coded as fuzzy variables (with four modalities each), we considered each of their modalities as independent traits in the PCA but associated them with a weight of 0.25 so that each original trait was weighted equally in the PCA. We combined both multivariate analyses with a co-inertia analysis (Dray et al., 2003). Such analysis allowed us to map communities, taxa and traits in the same multivariate space. We used the first two axes of the co-inertia analysis (66% of the covariance between traits and communities) as descriptors of the average trait structure of communities (e.g. Moretti & Legg, 2009).

We calculated multi-trait community functional diversity using the equivalent number of Rao’s quadratic entropy (Leinster & Cobbold, 2012). To integrate traits coded either continuously or as fuzzy variables, we calculated Gower’s distance. For each community, null functional diversity distributions were generated by permuting the columns (taxa) of the site-by-taxa abundance matrix. We then computed the standard effect sizes (SES) to evaluate the deviations of the observed functional metrics from the random expectation. SES are calculated as the observed metric value minus the mean of the null functional diversity values divided by the standard deviation of the null functional diversity values. A negative SES value indicates that the metric is lower than expected according to the null hypothesis. It can be interpreted as abiotic filtering being the main process structuring communities. Conversely, a positive SES value indicates that the metric is higher than expected. It can be interpreted as biotic interactions being the main process structuring communities. Finally a SES value close to 0 indicates that stochastic processes (e.g. colonization and extinction events) are the main processes structuring communities (Spasojevic et al., 2014).

#### Community structure modeling

Using linear mixed models, we related community statistics at each reach (the two co-inertia axes, taxa richness and functional diversity SES) to multiple predictors:

- Spatio-temporal connectivity of the community within the river network during the 30 days prior to sampling.
- Drying frequency and drying duration in the ten years before sampling.
- Day of the year (starting in 01-01-2021) as a degree 2 polynomial covariate to model the seasonal dynamic of the community. The variable was scaled.
- The identity of the drying river network to account for biogeographical differences among DRNs.
- A reach-level random effect to account for repeated sampling at the same reach and to account for spatial variation in environmental conditions among reaches (see Supplementary materials).
- Interaction terms between drying metrics and connectivity or day of the year to assess how intermittence interacts with spatio-temporal connectivity to shape community structure and how it may change the seasonal dynamic of communities.

For each community metric, we evaluated all the possible linear mixed models including all or a subset of terms, up to eight fixed effect terms using the function ‘dredge’ from R-package ‘MuMIn’. Models were ranked according to corrected Akaike information criterion (AICc). We retained the single model with the smallest AICc as the ‘best’ model, computed its marginal and conditional R^2^ (Nakagawa & Schielzeth, 2013) to access the variance explained respectively by fixed effects only and both fixed and random effects.

We ran a sensitivity analysis on the effect of the time windows chosen to compute drying frequency and average duration (60 days, 90 days, one year, two years, 5 years and 10 years). It showed that for a time window at least equal to one year, the models have similar high explanatory power (see Figure S3).

To better interpret diversity patterns, we ran a similar analysis on single trait community weighted means. Finally, we also ran alternative, more descriptive models on community metrics that use intermittence regime (categorical: perennial vs. intermittent) and DRN identity as covariates. Results are in the supplementary materials.

All analyses were conducted using R version 4.3.1 (R Core Development Team 2012) using the packages: ‘ade4’ (Dray & Dufour, 2007), ‘gawdis’ (de Bello et al., 2021), ‘lme4’ (Bates et al., 2014) and ‘MuMIn’ (Bartoń, 2012).

## Results

### Functional structure of perennial and intermittent reaches

The primary axis of the co-inertia analysis (48% of total inertia) between community and trait turnover was underpinned by a fecundity-longevity-dispersal trade-off (Figure 2a). Taxa with negative scores were often NI or OCH taxa and tend to display a long-lived adult phase, low fecundity, small body size, were overall less mobile (all dispersal traits have negative scores) and were more likely to exhibit a cocoon resistance form. Taxa with positive scores were often EPT taxa and other insect taxa (OI) and tend to display opposite trait features as well as an egg-based resistance form.

**Figure 2.**
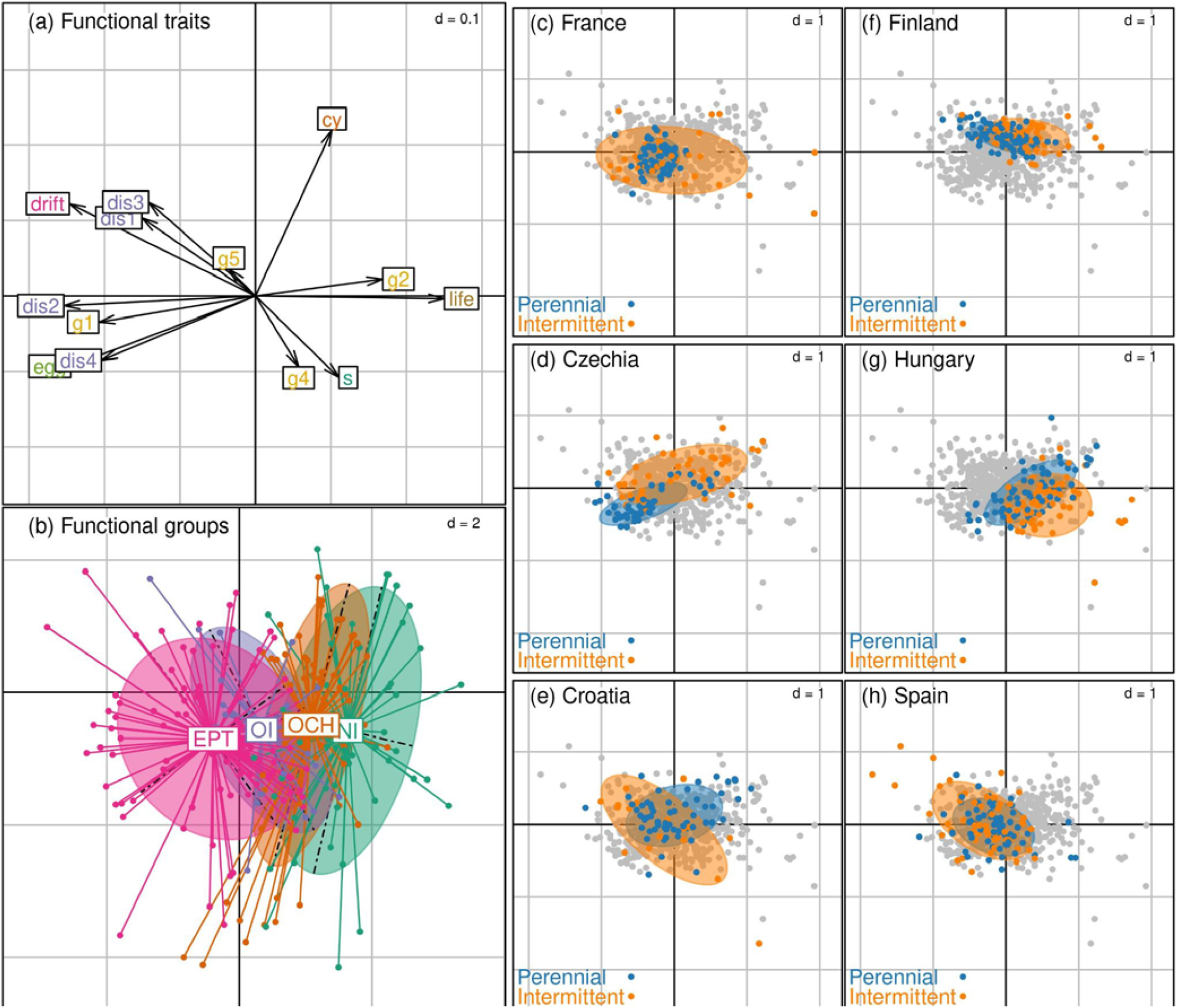
Overview of the functional structure of the river communities. All sub-panels represent the scores of communities, macroinvertebrate taxa and traits along the first two axes of the co-inertia analysis on community and trait data. (a) Scores of functional traits. The font color refers to each of the traits: the number of reproductive cycle (orange, ‘cy’), dispersal mode (purple: ‘dis1’ - passive aquatic; ‘dis2’ - active aquatic; ‘dis3’ - passive aerial; ‘dis4’ - active aerial), drift frequency (pink; ‘drift’); lifelong fecundity (green; ‘egg’); resistance form (yellow; ‘g1’ - Eggs, statoblasts, gemmules; ‘g2’ - Cocoons; ‘g4’ - Dormancy; ‘g5’ - None); adult life span (brown; ‘life’); body size (blue, “s”). (b) Scores of taxa grouped by taxonomic groups: EPT (pink) groups Ephemeroptera, Plecoptera and Trichoptera; OCH (orange) groups Odonata, Coleoptera and Hemiptera; OI (blue) groups other insect clades (i.e. Diptera and Megaloptera taxa); NI (green) groups all non insect clades. (c-h) Positions of sampled reach communities. In all subpanels, communities across all DRNs and sampling campaigns are displayed. In each subpanel, we highlighted the communities associated with each DRN in blue (perennial reach communities) and orange (intermittent reach communities), communities from other DRNs are in gray.

The second axis (17% of total inertia) separated taxa according to their reproductive strategy and some aspects of dispersal and resistance. Taxa with negative scores tend to have a single reproductive cycle per year, have a large body size and exhibit a dormancy resistance form. Taxa with positive scores tend to have opposite trait features, as well as a higher probability of drifting and having a passive aerial or aquatic dispersal mode. This axis was unrelated to the taxonomic groups (Figure 2b).

Community scores along the first two axes depended on their associated DRN and intermittence regime (Figure 2 c-h, Table S2). Perennial communities had a less variable position along the first two co-inertia axes than intermittent communities. (Ratio of variance of the first axis = 0.52, p-value < 0.001; second axis = 0.63, p-value < 0.001). This variability was due to a DRN-dependent position of intermittent vs. perennial reaches communities (Figure 2, Table S2).

Along the first co-inertia axis, perennial and intermittent communities in the Spanish, Croatian DRN had similar scores. Intermittent communities in the French, Czech, Finnish and Hungarian DRN had more negative scores than perennial communities. Along the second co-inertia axis, perennial and intermittent communities tend to have similar structure except for the Czech DRN where perennial communities had more negative scores than intermittent ones.

### Drivers of community structure

Linear models linking community metrics to drying duration and frequency, spatio-temporal connectivity, seasonality and biogeography explained a moderate proportion of the community metrics (marginal R^2^ between 0.13 and 0.33, Table 1). Linear models of community weighted means for all single traits but ‘number of reproductive cycles’ displayed a similar performance with a marginal R^2^ ranging from 0.20 to 0.38. In contrast, the number of reproductive cycles was not well explained with a marginal R^2^ of 0.06 (Table S3). Accounting for random effects (aka. site effects), the explained variance was much higher and ranged from 0.47 to 0.76 indicating that much of the variation of community metrics among sites remained unexplained by the fixed effects of our models.

**Table 1.** Summary of the linear mixed models between community metrics and intermittency, seasonal and connectivity factors. The default modality of the DRN factor referred to the Croatian DRN. DOY refers to ‘day of the year’; ‘Connect.’ to spatio-temporal connectivity; “Drying dur.” to drying duration; “drying freq.” to drying frequency.

| *Predictors* | Total density (log ind/m<sup>2</sup>) |  | Richness |  | 1st Coinertia axis |  | 2nd Coinertia axis |  | Func. diversity |  |
| --- | --- | --- | --- | --- | --- | --- | --- | --- | --- | --- |
|  | *Estimates* | *p* | *Estimates* | *p* | *Estimates* | *p* | *Estimates* | *p* | *Estimates* | *p* |
| (Intercept) | 7.26 | **<0.001** | 24.29 | **<0.001** | -0.05 | 0.684 | 0.05 | 0.380 | 0.91 | **<0.001** |
| DRN [Czechia] | 0.64 | **0.009** | 4.97 | **0.039** | -0.09 | 0.501 | -0.13 | 0.090 | -0.25 | 0.211 |
| DRN [Finland] | 0.04 | 0.874 | 0.44 | 0.851 | 0.17 | 0.202 | 0.06 | 0.402 | -0.19 | 0.334 |
| DRN [France] | -0.03 | 0.909 | 7.35 | **0.002** | -0.06 | 0.657 | -0.19 | **0.009** | 0.45 | **0.023** |
| DRN [Hungary] | 0.83 | **0.001** | 5.27 | **0.026** | 0.48 | **0.001** | -0.24 | **0.001** | 0.41 | **0.038** |
| DRN [Spain] | -0.58 | **0.018** | 7.35 | **0.002** | -0.30 | **0.036** | -0.01 | 0.906 | 0.08 | 0.697 |
| DOY | 4.61 | **<0.001** | 7.35 | 0.328 | 1.49 | **0.001** | -0.46 | 0.187 | -0.99 | 0.269 |
| DOY:Drying dur. |  |  |  |  |  |  | 0.17 | 0.124 |  |  |
| DOY:Drying freq. |  |  |  |  | -0.28 | 0.420 | -0.51 | 0.109 | 0.05 | 0.935 |
| DOY<sup>2</sup> | -7.47 | **<0.001** | -21.28 | **0.009** | -1.57 | **0.003** | -2.08 | **<0.001** | 1.59 | 0.121 |
| DOY<sup>2</sup>:Drying dur. |  |  |  |  |  |  | 0.36 | **0.003** |  |  |
| DOY<sup>2</sup>:Drying freq. |  |  |  |  | 1.05 | **0.007** | -0.79 | **0.027** | -1.98 | **0.008** |
| Connect. | 0.35 | 0.091 | 1.92 | 0.154 | -0.05 | 0.424 | 0.07 | 0.054 | 0.15 | 0.105 |
| Connect.:Drying dur. |  |  |  |  | -0.05 | **0.004** |  |  |  |  |
| Connect.:Drying freq. | 0.58 | **0.001** | 2.28 | 0.051 |  |  |  |  |  |  |
| Drying dur. | 0.07 | 0.085 |  |  | -0.00 | 0.971 | -0.01 | 0.419 |  |  |
| Drying freq. | -0.75 | **<0.001** | -6.61 | **<0.001** | 0.13 | **0.020** | 0.03 | 0.330 | -0.16 | **0.014** |
| **Random Effects** |  |  |  |  |  |  |  |  |  |  |
| $\sigma^2$ | 0.95 | | 38.47 | | 0.07 | | 0.04 | | 0.28 | |
| $\tau_{00}$ | 0.27 site | | 38.45 site | | 0.14 site | | 0.04 site | | 0.26 site | |
| ICC | 0.22 |  | 0.50 |  | 0.67 |  | 0.49 |  | 0.48 |  |
| N | 126 site |  | 126 site |  | 126 site |  | 126 site |  | 126 site |  |
| Observations | 634 |  | 638 |  | 636 |  | 636 |  | 636 |  |
| Marginal R<sup>2</sup> / Conditional R<sup>2</sup> | 0.332 / 0.479 |  | 0.191 / 0.595 |  | 0.264 / 0.759 |  | 0.235 / 0.607 |  | 0.131 / 0.544 |  |

#### Effect of flow intermittence on community structure

Spatio-temporal connectivity had no effect on the main community metrics when drying duration and drying frequency were close to 0 (Table 1). Highly connected communities displayed more individuals from taxa with a high probability of drifting (p < 0.001), a high score of active aquatic dispersal (p < 0.001) and passive aerial dispersal (p = 0.013).

Drying frequency had a substantial impact on most community structure metrics (Table 1). The total density of individuals, taxa richness and functional diversity decreased with drying frequency. The score of communities along the first co-inertia axis increased with drying frequency, indicating a higher proportion of OCH and NI individuals in more intermittent reaches (Figure 3, Table 1). Single trait analyses (Table S3-S5) indicated that this shift was due to a decrease in aquatic dispersal ability (active/passive) and an increase in average longevity. Taxa were also less likely to exhibit an egg-based resistance form and more likely to exhibit a cocoon-based resistance form.

**Figure 3.**
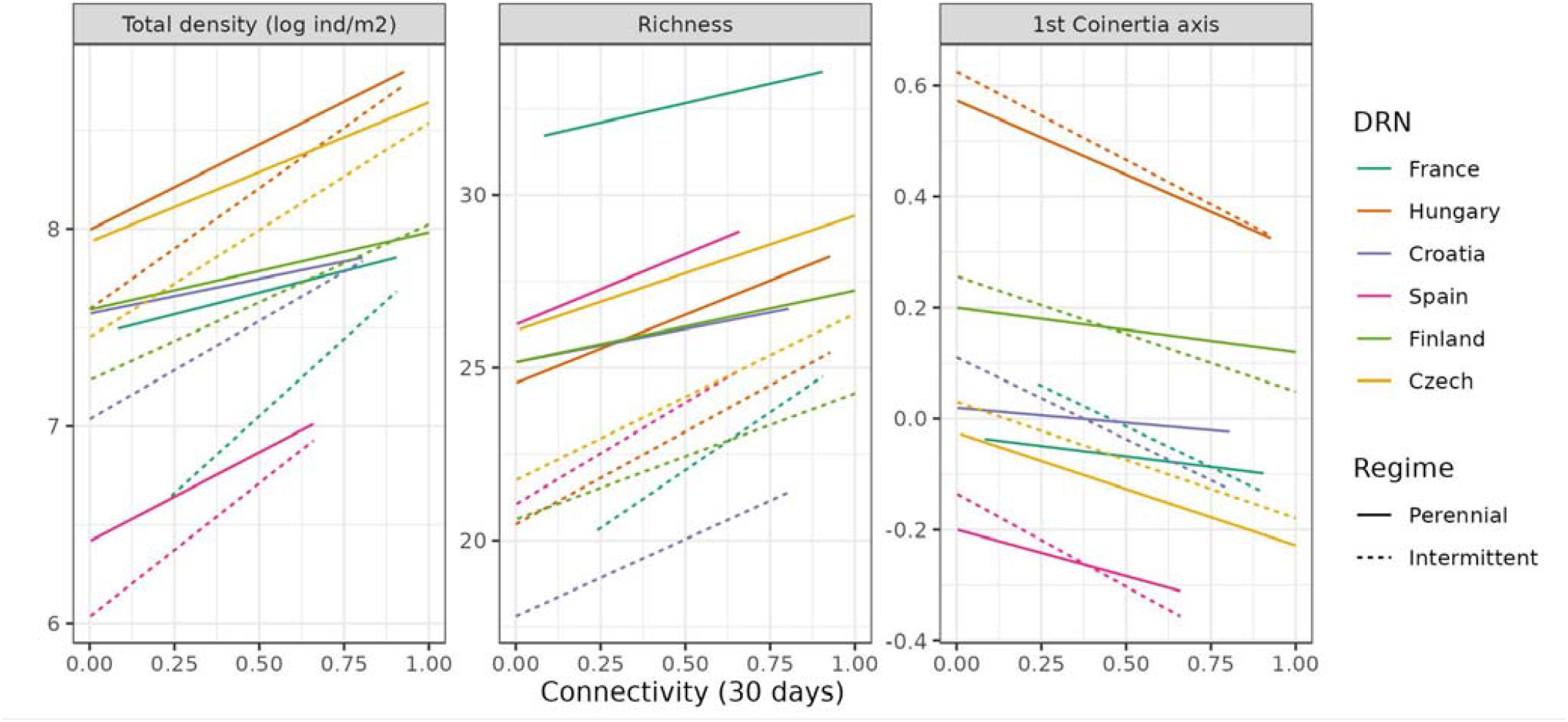
Effect of spatio-temporal connectivity on community structure. The graphics display the marginal effects of the spatio-temporal connectivity in the 30 days prior to sampling on community structure. To visualize the interaction effects of spatio-temporal connectivity, drying predictors and DRN, we displayed the relationships between community metrics and spatio-temporal connectivity for average levels of drying duration and drying frequency of perennial and intermittent reaches in each drying river network. Only community metrics that are significantly impacted by connectivity are displayed : total density of individuals (log-transformed), community richness, the first axis of the co-inertia analysis describing the main functional trait turnover between communities.

In communities with a high drying duration, taxa were more likely to have a dormancy-based resistance form (p< 0.001) and less likely to have a cocoon-based resistance form (p< 0.001) and less likely to have no resistance form (p=0.035).

Spatio-temporal connectivity significantly modulated the impact of intermittence on communities (Table 1, Figure 3). The negative impact of drying frequency on total density and (marginally) taxa richness was compensated when the reach was highly connected to other reaches. There also was a significant interaction between drying duration and connectivity for the first axis of co-inertia analysis (Table 1, Figure 3) indicating that highly connected communities with long drying duration had more EPT and OI individuals (Figure 1). Single trait analyses (Table S3-S5) indicated that this shift was essentially due to an increase in fecundity (p = 0.017), a decrease in the longevity (p = 0.012), an increase in aerial active dispersal ability (p = 0.003).

#### Seasonal variation of community structure

River community structure varied significantly across the year and this variation depended on the flow regime. There were significant bell-shaped relationships between day of the year and all community metrics (Table 1, Figure 4). Total density of individuals and taxa richness peaked in late spring to summer and communities also had higher scores along the first and second co-inertia axes indicating a higher proportion of OCH and NI taxa (compared to EPT and OI). Single trait analyses (Table S3-S5) indicated that these shifts were due to an increase in abundance of taxa with a high number of reproduction cycles, lower fecundity, higher longevity or larger body size.

**Figure 4.**
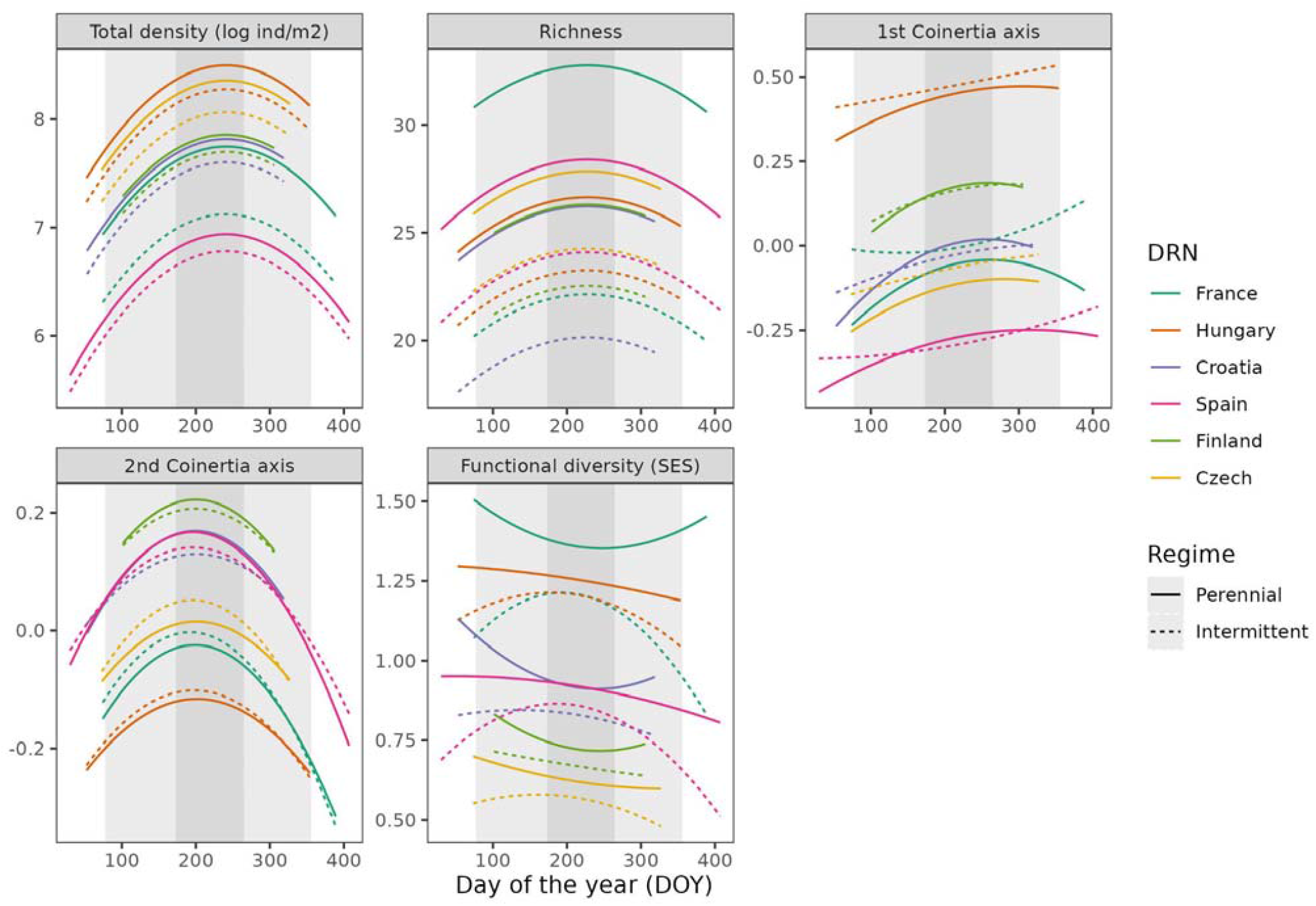
Seasonal variation of community structure. The graphics display the marginal effect of the day of the year (DOY) for the drying duration and drying frequency of perennial and intermittent river reaches in each drying river network. Community structure is described by the total density of individuals (log-transformed), community richness, the two first axes of the co-inertia analysis describing the main functional trait turnover between communities and the standard effect size of the functional diversity. Background areas indicate winter (white), spring and fall (light gray) and summer (gray) according to solstices and equinoxes dates.

Intermittence significantly impacted the seasonal variation of communities (Table 1, Figure 4). In particular, there was a significant interaction between drying frequency and day of the year for the first axis of the co-inertia analysis. It indicated that communities from reaches with a high drying frequency had a higher proportion of OCH and NI taxa (and low relative abundance of EPT and OI genera) after summer compared to perennial reaches. The functional diversity of perennial reaches did not vary across the year but peaked in late spring to summer in intermittent reaches (Table 1, Figure 4). Single trait analyses indicated that these shifts were likely due to taxa with a high fecundity and/or low longevity decreasing across the year in intermittent reaches whereas, they tended to increase in relative abundance after summer in perennial reaches.

#### Variation of community structure across DRNs

After accounting for differences in drying duration, drying frequency and connectivity, there was a significant effect of DRN on community structure.

Communities from the Hungarian DRN had a higher density of individuals (p < 0.001), richness (p = 0.002) and functional diversity (p = 0.036). They had a higher score on the first co-inertia axis indicating a higher proportion of OCH and NI, and a more negative score along the second co-inertia axis (Figure 2, Table 1). This was due to a higher proportion of taxa with a large body size, a low probability of drifting and low aquatic (passive and active) or aerial passive dispersal ability (Table S3, S4).

In contrast, communities from the Spanish DRN had a higher richness (p=0.002), a lower score along the first co-inertia axes reflecting a higher proportion of EPT and NI genera. This was due to a higher proportion of taxa with a short lifespan and a stronger aerial passive or active ability (Table S3-S4). The Spanish and Hungarian DRN communities both displayed a lower proportion of taxa with no resistance form compared to other DRNs (Table S5). However the dominant resistance strategy differed: egg-resistant form was more abundant in the Spanish DRN while cocoon-based or dormancy based resistance forms were more abundant in the Hungarian DRN (Table S5). Communities from the French DRN exhibited a higher taxa richness and functional diversity compared to the other DRNs. Like in the Hungarian DRN, communities scored lower on the second co-inertia axis (Figure 2) because of a higher proportion of large-bodied taxa.

## Discussion

A puzzling feature of intermittent river communities is that they show highly variable structure across space and time. Our spatio-temporal study of six drying river networks in Europe indicates that intermittent communities have highly variable taxonomic and functional structure across space and time compared to perennial communities, in accordance with other field studies (e.g. Crabot, Polášek, et al., 2021). Our study reveals how this variability is underpinned by differences in biogeographical location, drying regime, spatio-temporal connectivity and seasonal dynamics thus giving insights on how biogeography, disturbances, dispersal dynamics and environmental fluctuations interact with species traits to shape the structure of biological communities (Holyoak et al., 2020; Spasojevic et al., 2014; Tonkin et al., 2017).

### Drying frequency and duration have divergent effects on community structure

Our results indicate that flow intermittence, as a natural disturbance, deeply shapes the functional structure of macroinvertebrate communities and selects for taxa that exhibit specific traits to cope with drying. Consistent with previous studies (e.g. Arias-Real et al., 2021), high drying frequency communities had lower densities of individuals, taxa richness and functional diversity than their perennial counterparts, showing the important filtering exerted by recurring river drying events (Figure 3). More puzzling, our study also shows that high drying frequency vs. long drying duration lead to divergent community trait structure. High drying frequency selected for K-strategists with a weak dispersal ability and was associated with increased proportion of taxa using cocoons as a resistance form. Longer drying duration in combination with high spatio-temporal connectivity selected taxa with r-strategies (short lifespan, high fecundity) and strong dispersal abilities. Interestingly drying frequency and duration were associated respectively to cocoon resistance form and desiccation-resistant egg forms or dormancy showing that different aspects of flow intermittence select for different resistance forms in addition to different life-history strategies.

A possible explanation for this pattern is that during frequent but short drying events the riverbed may not dry severely and can provide refugia in the hyporheic zone for macroinvertebrates with adequate resistance adaptations (Viza et al., 2023). However, long drying duration may favor resilience-based strategies by driving the community beyond a critical ecological threshold (see Aspin et al., 2019) where resistance strategies are not sufficient to cope with drying. This may explain contrasting results in experimental rewetting experiments across river networks (Datry et al., 2012; Folch de la Iglesia et al., 2020); as well as the high proportion of resilience strategies in intermittent stream communities from drier climates, including mediterranean and semi-arid (Giam et al., 2017; Pineda-Morante et al., 2022).

### Connectivity shifts the structure of intermittent river communities

A key question regarding connectivity and community structure is whether dispersal processes align more with a deterministic or stochastic perspective on meta-community dynamics. Our study provides a complex answer. On one hand, spatio-temporal connectivity had a positive effect on the total density of individuals and taxa richness, but only a positive non-significant, effect on functional diversity (Table 1), giving limited support to the hypothesis that increased connectivity would increase stochasticity and dilute the impact of deterministic processes by increasing the abundance of poorly adapted species, and thus community density and richness (Spasojevic et al., 2014). On the other hand, our study shows that increased spatio-temporal connectivity is associated with a higher proportion of r-strategists with high dispersal ability in intermittent communities but not in perennial communities (Figure 2). This supports the idea that meta-community dynamics in DRNs are trait-based, suggesting that patch dynamics drive assembly, with good dispersers thriving in disturbed segments if connected to perennial ones (Cañedo□Argüelles et al., 2015; Sarremejane, Mykrä, et al., 2017).

In addition, our analysis indicates that spatio-temporal connectivity positively influences the relative abundance of taxa with an egg-based resistant form. Thus, the idea that connectivity does favor resilient strategies over resistance strategies in intermittent reaches was not as clear-cut as hypothesized. In particular, egg-based resistance was related to life-history traits related to resilience strategies across macroinvertebrate taxa (fecundity, r = 0.41, p<0.001; longevity, r = −0.31, p<0.001); this indicates that coordinated resilience and resistance strategies over the life cycle of some taxa may be necessary for them to thrive in intermittent segments.

### Intermittence modifies the seasonal dynamic of communities

In general, the seasonal variation of communities is generally explained by increased population growth and resource depletion in summer, leading to higher density-dependence and a peak in community diversity (Mellard et al., 2019). Our study shows that DRN macroinvertebrate communities conform to this general pattern (Figure 4). We further provide insights into how species respond to seasonal variations through their functional traits and how natural disturbance caused by flow intermittence modifies the seasonal variation of macroinvertebrate communities.

Perennial reaches were characterized by similar assemblages in spring and autumn composed of r-strategists and a summer assemblage composed of K-strategists. This is consistent with the idea that r-strategies dominate both in spring and autumn during rewetting, while slow-growing, competitive taxa prevail in summer due to resource depletion, a pattern observed in other ecosystems (e.g. terrestrial plants, Doležal et al., 2019). Despite this functional shift, there was no significant seasonal variation of the functional diversity of perennial segments (Table 1) suggesting that assembly processes remain constant throughout the year in perennial reaches. In addition to the r-to K-strategist shift, we found that multi-voltism and small body size are prevalent traits for active summer taxa (Figure 4). Multi-voltism, influenced by high temperatures (Altermatt, 2009) and seasonal variations of temperature (Bonacina et al., 2023), appears to be a key trait in the seasonal variation of macroinvertebrate communities.

However, the seasonal dynamic was different in intermittent reaches. Communities shifted from being dominated by r-strategists in spring to being dominated by K-strategists in both summer and autumn (Figure 4). This distinct “autumn assemblage” was further less taxa-rich than the summer assemblage. Functional diversity peaked in summer suggesting that biotic interactions structure more communities in summer than in spring and autumn. Conversely functional diversities were more likely to be random or convergent in spring and autumn. This may indicate that stochastic processes are more important in spring and autumn during rewetting: as intermittent streams reconnect to the rest of the drying river network (Figure S4), community assembly in intermittent communities may be less structured by deterministic processes and more by random colonization and extinction events.

### Community trait structure depends on the biogeographical and environmental context

Beyond differences in intermittence regime and connectivity dynamics, our work suggests that macroinvertebrate community assembly differs among DRNs. Fitting our expectations, after accounting for intermittence and connectivity, the Mediterranean communities of the Spanish DRN were on average more taxa-rich and had a higher proportion of taxa with resilience strategies. However, biogeographical differences among DRN communities were more prominent between non-mountainous DRNs (Hungary and Finland) and mountainous DRNs (Figure 1, Table 1). This suggests that in our study the potential effect of the evolutionary history of intermittent river biodiversity associated with the Mediterranean vs. temperate regions (Bonada et al., 2007) is obscured by the geomorphological context associated with each DRN. Non mountainous river networks, characterized by slow flow velocity and small particle size in the riverbed (Figure S2), harbored distinct community structures with more K-strategists (often belonging to OCH and NI) and a lower proportion of taxa with strong dispersal abilities. In contrast, communities from mountainous DRNs (Spain, France, Croatia, and Czechia) contained more r-strategists (often belonging to EPT), and exhibited a more resilient profile with more fecund and/or mobile taxa and a higher proportion of good dispersers.

Additionally, our study offers new insights into the determinants of resistance strategies across river networks. Communities from the Spanish and Hungarian DRNs shared a similar low proportion of taxa without any resistance strategy (Table S5), despite being the most dissimilar DRNs in terms of environmental conditions (Figure S1) and overall community structure (Figure 1). Our findings suggest that different resistance strategies may be optimal depending on macroinvertebrate r/K strategies and the local environmental context of DRNs. For example, resistance strategies based on eggs may be more optimal for resilient r-strategists typically found in mountainous and/or Mediterranean areas. Conversely, resistance strategies based on cocoons or dormancy may be more suitable for non mountainous, low-discharge rivers that favor K-strategists with a weak dispersal ability.

## Conclusion

Our study answers recent calls to better assess the joint influence of natural disturbances, connectivity on the seasonal dynamics of meta-communities (Holyoak et al., 2020; Tonkin et al., 2017). There are three key take-away messages:

First, a striking result of our study is how different facets of natural disturbance regime, here duration, frequency and their impact on connectivity dynamics among communities select for different ecological strategies and lead to divergent community structure. This calls for an explicit distinction between these when studying biological communities. Furthermore, as future climate and increased anthropic pressures on water resources should increase river network intermittence and fragmentation (Döll & Schmied, 2012; Zipper et al., 2021), incorporating connectivity measures into conservation planning may be essential to preserve river network structure and the natural habitat heterogeneity generated by it to ensure community recovery after drying.

Second, we showed how the respective influence of stochastic vs. deterministic assembly processes vary across the year and lead to divergent seasonal dynamics between non-disturbed and disturbed communities. As global change is expected to modify climate seasonality and thus disrupt the seasonal aspect of community assembly (Pau et al., 2011), better assessing the temporality of biological communities and how it is determined by species traits is an essential task for community ecologists.

Finally, our study shed some light on the complex patterns linked to life history and resilience/resistance traits in regards to different biogeographical and environmental contexts. Recognizing the contingency among macroinvertebrate traits may enhance our understanding of how different ecological strategies are selected in DRNs and may further call for differentiated river management strategies according to the local regional context.

## Data availability statement

All data and the R-script underlying this study are accessible on *https://figshare.com/s/8824ca412357d71ec569*

## Conflict of interest statement

The authors declare that they have no conflicts of interest.

## Notes

### Competing Interest Statement

The authors have declared no competing interest.

