## Supplementary materials for "Natural disturbances and connectivity shape the seasonal variability of aquatic macroinvertebrate communities across Europe"

### Supplementary material 1: extended methods

**Community sampling** - Quantitative samples were collected with a Surber, Hess, or modified Surber (mesh size 500 µm) sampler and the same device was used in all reaches and campaigns within each DRN. At each reach, the subsamples were taken proportionally from different habitats available (hard substrate, sand and silt, dead organic material, vegetated river edges, macrophytes, and mosses). The total sampled area depended on the mean stream or pool width (Flowing: width <3 m = 0.5 m^2^, width 3-10 m = 1 m^2^, and width >10 m = 1.5 m^2^. Only pools: width <1 m = 0.1 m^2^, width 1-5 m = 0.3 m^2^, and width >5 m = 0.5 m^2^). Subsamples were sieved (10 mm, 2 mm, and 0.5 mm sieves) to remove large organic particles from the samples and pooled into a composite sample. The samples were preserved in 96 % ethanol and later identified as often as possible to genus level, except for younger larval instars which were identified to the lowest possible taxonomic level and Diptera to family level. In case identification of higher taxonomic group (HTG) was due to small/damaged individuals, these were separated into the genera at that site. We excluded 886 individuals belonging to terrestrial fauna (terrestrial Coleoptera and Isopoda taxa) or to the meiofauna (belonging to the Nematoda and Nematomorpha phylums and the Hydrachnidae family). In case identification of higher taxonomic group (HTG) was due to small or damaged individuals, these were separated into the genera at that site.

Separation was done as described here:

A) In case all the possible genera were found at that specific sampling occasion (Sample ID), then the numbers of individuals (NOI) of the HTG were separated and added to the existing genera in the ratio of the NOI of these genera at that specific sampling occasion.

B) In case not all possible genera were found at that specific sampling occasion (Sample ID), then the (NOI) of the HTG was separated and added to the existing genera plus all the other possible genera at that sampling site (summed for all sampling occasions) in the ratio of the NOI of these genera at that sampling site (summed for all occasions).

B.2) In some cases, based on experts opinion, other or not just one sampling site's NOI ratio was used, but several, if these sites were close/similar.

If none of the above was possible, or the HTG identification was due to other reasons (e.g. it was not possible to identify some taxa) then the HTGs were kept as they were.

Also, in cases where the lower taxonomic resolution could be reached only episodically, and these identifications were not consistent, then these Genere were moved back to the HTG identification (i.e. some genera were changed back to Family level).

**Trait data** - Trait data were extracted from two overlapping genus-level databases (Sarremejane et al., 2020; Tachet et al., 2010) that coded traits as fuzzy variables. We selected life-history traits and known resistance and resilience traits based on the overview provided by Stubbington et al., (2017): maximum body size in cm (≤ 0.25, ]0.25, 0.5], ]0.5, 1], ]1,2], ]2,4], ]4, 8], > 8]), dispersal mode (passive aquatic dispersal, active aquatic dispersal, passive aerial dispersal, active aquatic dispersal); drift probability (rare, occasional, frequent); longevity (< one week; one week to one month; one month to one year; over one year); fecundity as lifelong number of eggs per female (<100; 100-1000; 1000-3000; > 3000); number of reproductive cycles per year (<1; 1; >1); resistance forms (eggs; cocoons; diapause/dormancy; none). Each modality is associated with a weight for each taxa. When trait data was available in both databases, we used preferentially the trait data from Sarremejane et al. (2020). A total of 8.9 x 10^5^ individuals belonging to 223 taxa were associated with genus-level trait data. For sampled individuals identified at family level (~ 4.5 x 10^5^ individuals classified in 35 taxa), we averaged the weights for all the genera recorded in the trait database that belonged to their respective family.

For traits with ordered modalities (maximum body size, longevity, lifelong fecundity, drift, number of reproductive cycles per year), we created ‘continuous traits’. For each taxon, we ranked the modalities as integer numbers and calculated the weighted average of that rank. Missing traits values for traits were imputed with chained random forests using the R-package ‘missRanger’ (Mayer, 2019). 208 out of 290 taxa had no missing trait values before imputation.

For the trait-based analyses, we excluded taxa that meet the following criteria: (1) taxonomic assignment was coarser than family-level. (2) There were missing values for four or more of the seven traits before trait imputation. This led to the exclusion of 25 taxa representing 2 842 individuals. The resulting loss in total number of individuals was minimal enough across community samples (median 0.04%, maximum 25%) and we decided to not exclude any community samples from the analysis. In total, across 638 community samples, we sorted and identified over 1.35 millions aquatic macroinvertebrate individuals belonging to 290 taxa (240 at the genus level, 45 at the family and 6 at a higher taxonomic level). For trait-based analyses, we analyzed 368 community samples containing a total of 1.3 x 10^6^ individuals classified in 265 taxons.

### Supplementary material 2: environmental variations across communities

At each sampling location and time, we measured a set of environmental variables reflecting the reach geomorphological, hydrological and physico-chemical characteristics (Figure S1, S2, Table S1).

To evaluate how sampled reaches differ across DRNs, we computed a between-class analysis on the PCA of the table environmental variables (Dolédec & Chessel 1987); it revealed that DRNs differ mostly according to two main environmental gradients (48% of variance explained by the first axis and 25% by the second). The first axis separated the mountainous DRNs Croatia, France, Spain characterized by larger particle size (i.e. a high proportion of cobbles and boulders), higher elevation, pH and oxygen concentration, from the non-mountainous DRNs of Hungary and Finland that have smaller particle size (high proportion of silt in Hungarian rivers, and sand in Finnish DRN), a more important riparian cover. The second axis was related mostly to intermittence and opposed the Spanish and Hungarian DRN to the Finnish DRN. The other DRNs have a variable and intermediate position.

To evaluate how sampled reaches differ within DRNs, we computed a within-class analysis on the PCA of the environmental variables revealed that communities within DRNs differed mostly according to a first axis (21% of variance) linked to hydrological characteristics (river width, depth and velocity). It opposed communities sampled in perennial reaches vs. communities sampled in intermittent reaches (Figure S2, right). The second axis was more related to the seasonal dynamic of reaches and opposed communities sampled during the cold months compared to communities sampled in summer. The former tend to have a higher connectivity, oxygen concentration and lower temperature than the latter.

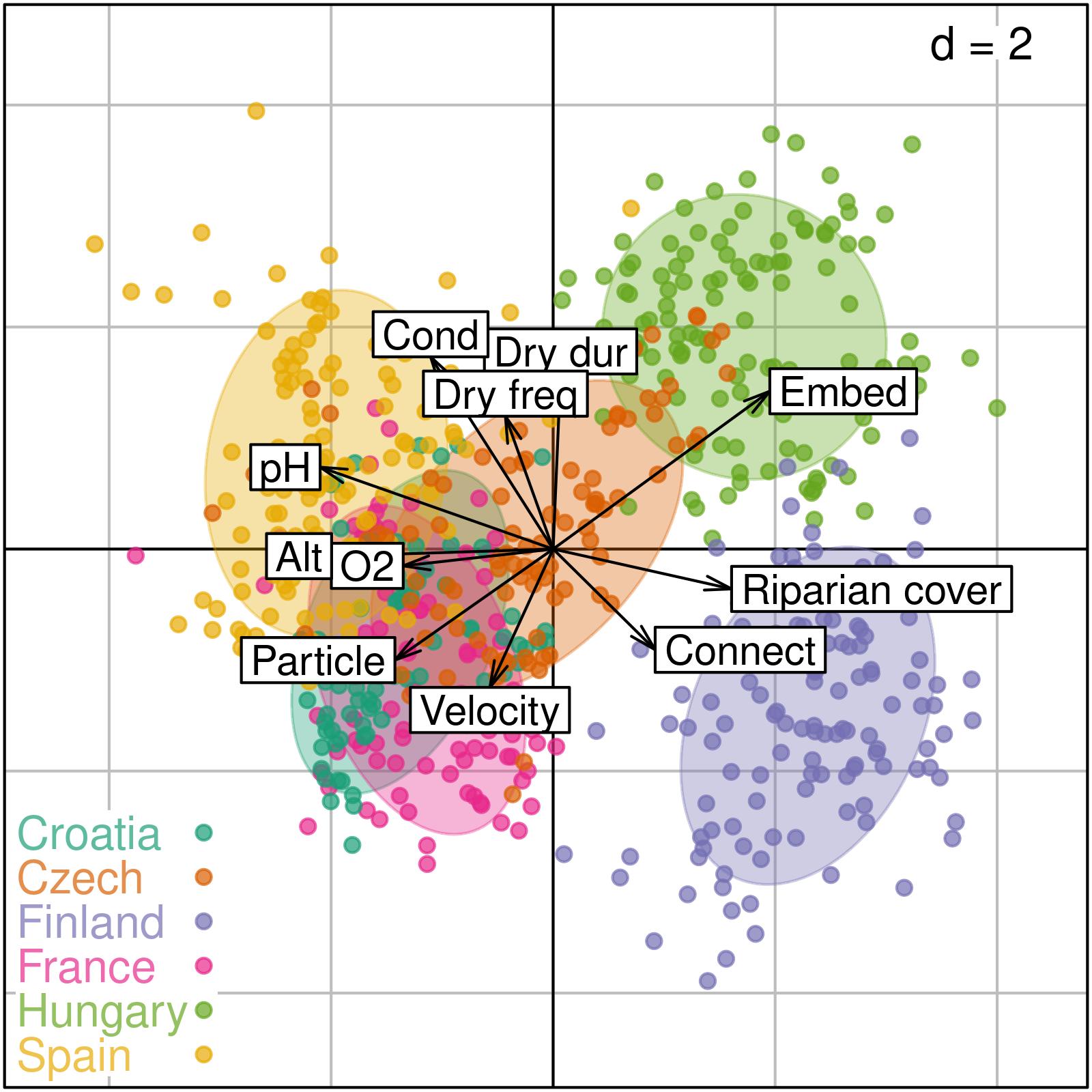

**Figure S1 - Environmental variation across drying river networks (DRN).** Colors and ellipses group sampled communities according to the DRN. Only the main environmental predictors are displayed (see Table S1 for a complete list with measurement details). Alt: altitude. Cond: conductivity. Connect. : spatio-temporal connectivity in the last 30 days. Dry dur: drying duration in the last 10 years (square root transformed). Dry freq: drying frequency in the last 10 years (square root transformed). Embed: embeddedness. O2: dioxygen concentration. Particle: riverbed particle size. pH: pH. Riparian cover : riparian vegetation cover. Velocity: average flow velocity (log transformed).

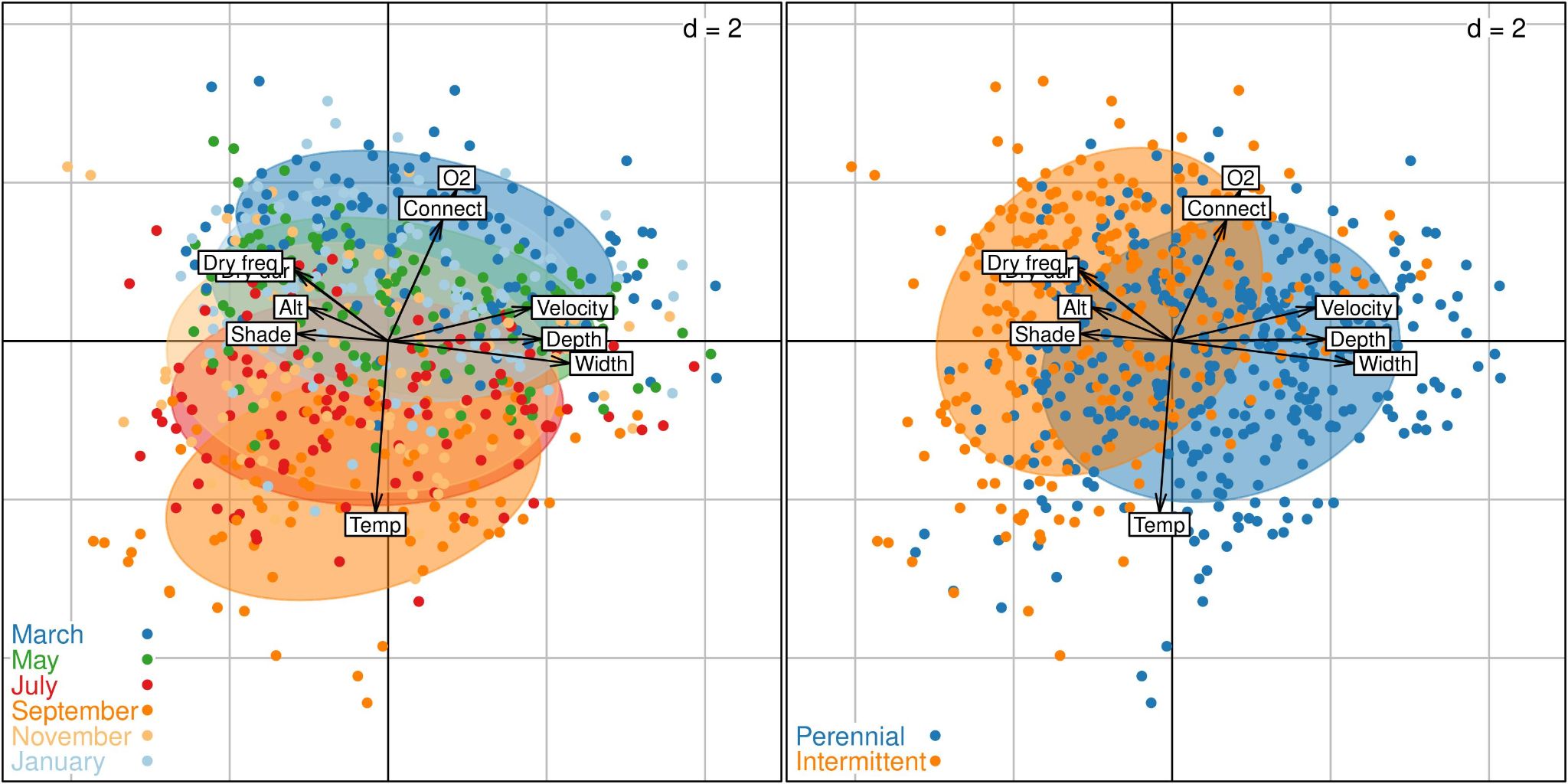

**Figure S2 - Environmental variation within drying river networks.** Colors and ellipses group sampled communities according to the month of sampling (left) and the intermittence regime (right). Only the main environmental predictors are displayed (see Table S1 for a complete list with measurement details). Alt: altitude. Connect. : spatio-temporal connectivity in the last 30 days. Depth: river depth (log transformed). Dry dur: drying duration in the last 10 years (square root transformed). Dry freq: drying frequency in the last 10 years (square root transformed). O2: dioxygen concentration. Particle: riverbed particle size. pH: pH. Shade: percentage of shade. Temp: water temperature. Velocity: average flow velocity (log transformed). Width: stream average wetted width (log transformed)

| Variable name | Units | Time sampled | Sampling location | Measurement details | Between-DRN PCA (Figure S1) | | Within DRN PCA (Figure S2) | |
| --- | --- | --- | --- | --- | --- | --- | --- | --- |
|  |  |  |  |  | 1st axis score | 2nd axis score | 1st axis score | 2nd axis score |
| Altitude | m | Once | One point in the reach | GIS | -0.67 | -0.02 | -0.34 | 0.14 |
| Slope | % | Once | Whole reach | Clinometer, a mean value along the whole reach | -0.33 | 0.12 | -0.25 | 0.05 |
| Average wetted width | m | Each time | Whole reach | Visual inspection of the whole reach or obtained from the measurements of the minimum and maximum wetted widths | -0.33 | -0.14 | 0.77 | -0.09 |
| Embeddedness (%) | 0-100% | Each time | Whole reach | Visual inspection of the whole reach | 0.65 | 0.47 | 0.01 | -0.08 |
| Particle size |  | Each time | Whole reach | Visual inspection of the whole reach | -0.47 | -0.33 | 0.01 | 0.13 |
| % riparian cover in the riparian area | 0-100% | Once, in spring | Whole reach | Visual inspection of the whole reach | 0.54 | -0.12 | -0.07 | -0.05 |
| % shade | 0-100% | Once, in spring | Whole reach | Visual inspection of the whole reach | 0.37 | -0.06 | -0.39 | 0.03 |
| Maximum depth | cm | Each time | One point in the reach | Rigid meter rule | -0.30 | -0.12 | 0.65 | 0.01 |
| Velocity | m/s | Each time | Every subsample of each sample | Velocimeter | 0.60 | 0.14 | 0.60 | 0.14 |
| Water temperature | ºC | Each time | One point in the reach | Thermometer | -0.24 | 0.07 | -0.05 | -0.72 |
| pH |  | Each time | One point in the reach | pH-meter | -0.70 | 0.25 | 0.10 | -0.15 |
| Conductivity | microS/cm | Each time | One point in the reach | Conductivity meter | 0.05 | -0.24 | 0.05 | -0.24 |
| Oxygen | mg/l | Each time | One point in the reach | Oxygen probe | -0.45 | -0.05 | 0.29 | 0.64 |
| Drying frequency | N/yr | 10 years | / | Hydrological modeling (see methods) | 0.02 | 0.52 | -0.45 | 0.33 |
| Drying duration | days | 10 years | / | Hydrological modeling (see methods) | -0.14 | 0.40 | -0.39 | 0.30 |
| Spatio-temporal connectivity | / | 30 days prior to sampling | / | Hydrological modeling (see methods) | 0.31 | -0.30 | 0.23 | 0.51 |

**Table S1 - Environmental variables measured in each sampled community.** We indicate measurement details and the score of those variables along the first two axes of the between-DRN PCA (Figure S1) and the within-DRN PCA (Figure S2).

### Supplementary material 3: Flow intermittence statistics across space, time and seasons

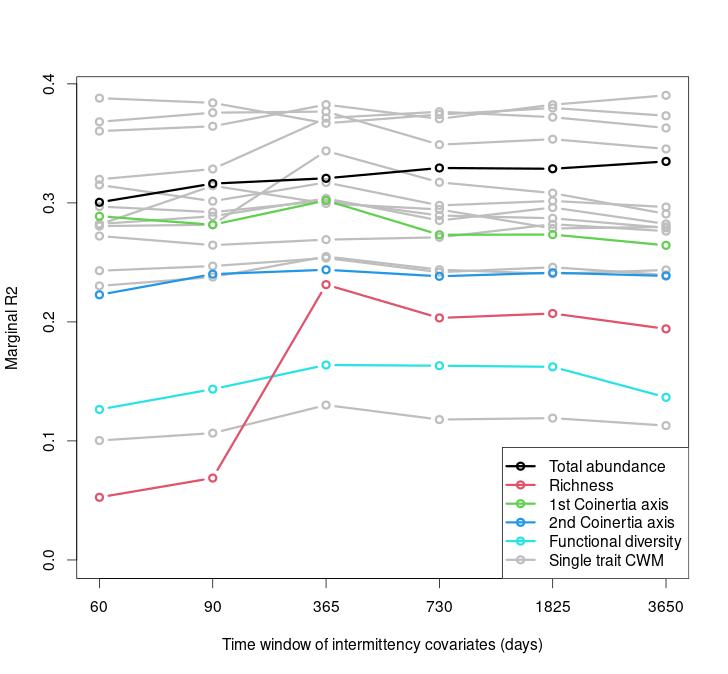

**Figure S3. Effect of the time window** used to calculate drying frequency and drying duration on the marginal R2 of the model.

We assessed the covariation of spatio-temporal connectivity, drying duration and drying frequency across the year with day of the year (starting in 01-01-2021), drying river network and reach type (intermittent vs. perennial). To do so, we used general additive models with day of the year as a smooth term and DRN and flow intermittence regime (perennial vs. intermittent) as categorical factors.
The different facets of flow intermittence varied strongly according to drying river networks, between perennial and intermittent reaches and across the year (Figure S4). Spatio-temporal connectivity varied across the year (F = 94.94, p < 0.001) with reaches being more connected in spring and autumn than in summer (Figure 1a). It varied across DRNs (F = 111.2, p< 0.001) and was highest in Finland compared to other DRNs (Figure 1b). Spatio-temporal connectivity was marginally lower in intermittent than in perennial reaches (F = 3.822, p = 0.051).
Since average drying duration and frequency were computed from modeled hydrological data, these metrics were not equal to zero in reaches classified as perennial from expert knowledge. Still as awaited, both drying frequency (F = 351.21, p < 0.001, Figure S4c) and drying duration (F = 325.68, p < 0.001, Figure 2d) were higher in intermittent than in perennial reaches.
Drying duration and drying frequency also varied across DRNs (F = 61.1, p < 0.001 and F = 43.09, p < 0.001, respectively) with intermittent reaches from the Spanish, Hungarian and Croatian DRN exhibiting the longest drying events and the highest drying frequencies (Figure S4d).
After averaging drying metrics by sampling reach, they were found to be linked by a degree 2 polynomial relationship (r^2^ = 0.56, p < 0.001, Figure S5) but that correlation was only moderate after excluding reaches with no drying event in the past ten years (r^2^ = 0.22, p = 0.010).

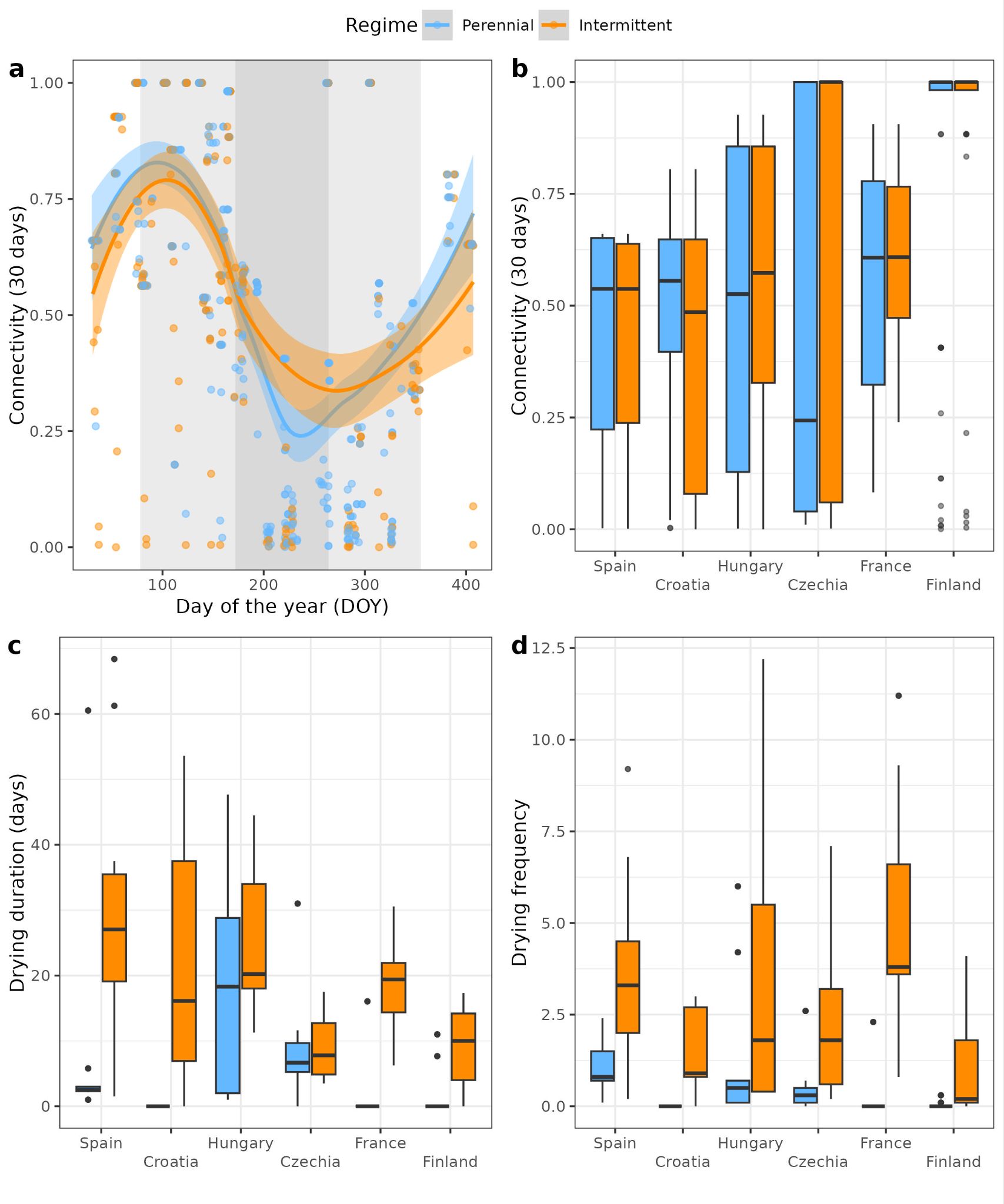

**Figure S4. Variation of connectivity, drying duration and drying frequency of studied river reaches according to stream type, the day of the year and drying river network.** We display the relationship between connectivity and day of the year (a) or the drying river network (b) as well as the variation of drying duration (c) and drying frequency (d) of sampled reaches across drying river networks. Background areas in subpanel (a) indicate winter (white), spring and fall (light gray) and summer (gray) according to solstices and equinoxes dates. DRN are arranged by latitude (south to north).

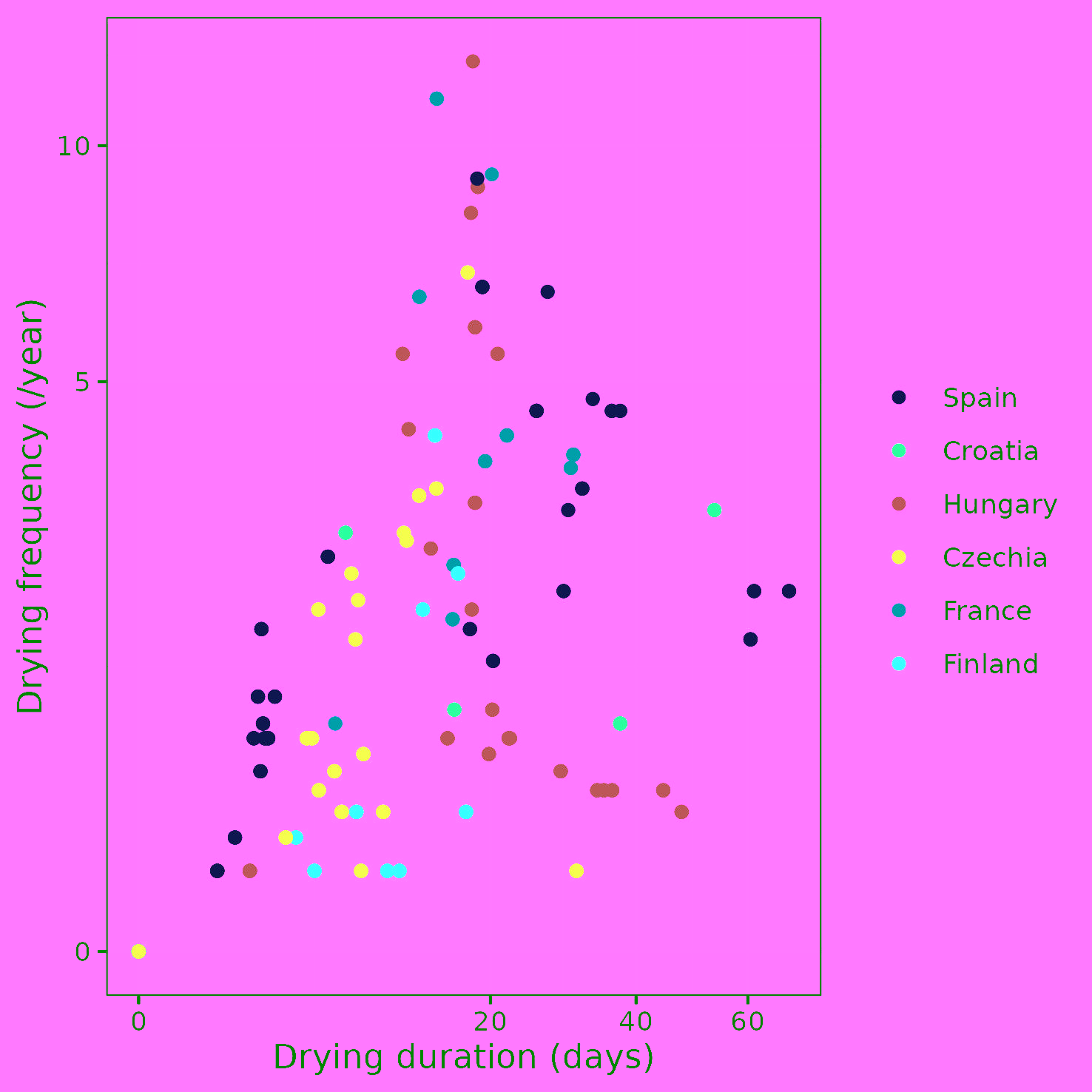

**Figure S5. Relationship between drying frequency and drying duration** (calculated on a time window of 10 years). Note that the (0,0) point overlay value pairs linked to perennial reaches.

### Supplementary material 4: alternative modeling of community structure

|  | **Total density (log ind/m2)** | | **Richness** | | **1st Coinertia axis** | | **2nd Coinertia axis** | | **Func. diversity** | |
| --- | --- | --- | --- | --- | --- | --- | --- | --- | --- | --- |
| *Predictors* | *Estimates* | *p* | *Estimates* | *p* | *Estimates* | *p* | *Estimates* | *p* | *Estimates* | *p* |
| (Intercept) | 7.78 | **<0.001** | 26.97 | **<0.001** | 0.01 | 0.957 | 0.13 | **0.027** | -0.75 | **<0.001** |
| Regime [Intermittent] | -1.47 | **<0.001** | -11.56 | **0.001** | -0.23 | 0.250 | -0.12 | 0.243 | 0.33 | 0.207 |
| DRN [Czechia] | 0.25 | 0.385 | 6.68 | **0.012** | -0.43 | **0.008** | -0.35 | **<0.001** | -0.57 | **0.006** |
| DRN [Finland] | -0.07 | 0.788 | 3.21 | 0.186 | -0.03 | 0.825 | 0.08 | 0.325 | -0.34 | 0.071 |
| DRN [France] | -0.01 | 0.966 | 4.92 | 0.052 | -0.23 | 0.136 | -0.21 | **0.009** | 0.22 | 0.259 |
| DRN [Hungary] | 0.52 | 0.055 | -1.42 | 0.573 | 0.33 | **0.031** | -0.20 | **0.011** | 0.60 | **0.002** |
| DRN [Spain] | -1.47 | **<0.001** | 1.45 | 0.566 | -0.16 | 0.287 | -0.18 | **0.028** | 0.17 | 0.388 |
| Regime:DRN [Czechia] | 1.35 | **0.005** | -4.22 | 0.322 | 0.78 | **0.003** | 0.51 | **<0.001** | 0.26 | 0.440 |
| Regime:DRN [Finland] | 1.72 | **<0.001** | -1.21 | 0.774 | 0.53 | **0.036** | 0.13 | 0.339 | 0.23 | 0.489 |
| Regime:DRN [France] | 0.04 | 0.935 | 1.74 | 0.686 | 0.56 | **0.031** | 0.06 | 0.640 | -0.21 | 0.529 |
| Regime:DRN [Hungary] | 0.84 | 0.066 | 7.56 | 0.065 | 0.46 | 0.061 | -0.06 | 0.650 | -0.90 | **0.006** |
| Regime:DRN [Spain] | 1.17 | **0.011** | 3.39 | 0.405 | 0.01 | 0.965 | 0.26 | 0.051 | -0.71 | **0.027** |
| **Random Effects** | | | | | | | | | | |
| σ^2^ | 1.02 | | 39.79 | | 0.09 | | 0.04 | | 0.26 | |
| τ00 | 0.21 site | | 26.56 site | | 0.11 site | | 0.03 site | | 0.16 site | |
| ICC | 0.17 | | 0.40 | | 0.56 | | 0.38 | | 0.38 | |
| N | 126 site | | 126 site | | 126 site | | 126 site | | 126 site | |
| Observations | 634 | | 638 | | 636 | | 636 | | 636 | |
| Marginal R^2^ / Conditional R^2^ | 0.324 / 0.440 | | 0.306 / 0.584 | | 0.306 / 0.696 | | 0.264 / 0.545 | | 0.195 / 0.504 | |

### Table S2. Summary of the linear mixed models between main community metrics and intermittency regime as a factor (perennial vs. intermittent) and DRN. The default modality of the DRN factor referred to the Croatian DRN.

### Supplementary material 5: single trait community weighted mean models

|  | **Number of repro. cycles per year** | | **Lifelong fecundity** | | **Adult life span** | | **Body size** | | **Propension to drift** | |
| --- | --- | --- | --- | --- | --- | --- | --- | --- | --- | --- |
| *Predictors* | *Coef* | *p* | *Coef* | *p* | *Coef* | *p* | *Coef* | *p* | *Coef* | *p* |
| (Intercept) | 2.30 | **<0.001** | 1.96 | **<0.001** | 2.32 | **<0.001** | 3.68 | **<0.001** | 1.68 | **<0.001** |
| DRN [Czech] |  |  | 0.13 | **0.009** | -0.11 | 0.157 | 0.06 | 0.248 | -0.03 | 0.435 |
| DRN [Finland] |  |  | -0.10 | **0.048** | -0.01 | 0.862 | -0.04 | 0.404 | -0.04 | 0.289 |
| DRN [France] |  |  | 0.11 | **0.034** | -0.03 | 0.660 | 0.13 | **0.007** | -0.04 | 0.380 |
| DRN [Hungary] |  |  | -0.07 | 0.201 | 0.19 | **0.018** | 0.09 | **0.038** | -0.24 | **<0.001** |
| DRN [Spain] |  |  | 0.03 | 0.498 | -0.33 | **<0.001** | -0.22 | **<0.001** | -0.07 | 0.081 |
| DOY | -0.08 | 0.305 | -0.30 | 0.111 | 1.32 | **<0.001** | 0.35 | 0.069 | -0.39 | **0.001** |
| DOY:Drying dur. |  |  | -0.15 | **0.018** |  |  |  |  |  |  |
| DOY:Drying freq. |  |  | 0.56 | **0.001** | -0.42 | 0.058 |  |  |  |  |
| DOY^2^ | -0.78 | **<0.001** | 1.08 | **<0.001** | -1.07 | **0.001** | 1.16 | **<0.001** | 0.12 | 0.382 |
| DOY^2^:Drying dur. |  |  | -0.09 | 0.201 |  |  |  |  |  |  |
| DOY^2^:Drying freq. |  |  | -0.46 | **0.016** | 0.77 | **0.002** |  |  |  |  |
| Connect. |  |  | 0.01 | 0.697 | 0.02 | 0.590 | -0.06 | **0.030** | 0.08 | **<0.001** |
| Connect.:Drying dur. |  |  | 0.02 | **0.017** | -0.03 | **0.012** |  |  |  |  |
| Drying dur. |  |  | 0.00 | 0.917 | -0.00 | 0.901 |  |  |  |  |
| Drying freq. |  |  | -0.03 | 0.131 | 0.06 | **0.049** |  |  |  |  |
| **Random Effects** | | | | | | | | | | |
| σ^2^ | 0.01 | | 0.01 | | 0.03 | | 0.03 | | 0.01 | |
| τ00 | 0.01 site | | 0.02 site | | 0.04 site | | 0.01 site | | 0.01 site | |
| ICC | 0.62 | | 0.63 | | 0.60 | | 0.34 | | 0.55 | |
| N | 126 site | | 126 site | | 126 site | | 126 site | | 126 site | |
| Observations | 636 | | 636 | | 636 | | 636 | | 636 | |
| Marginal R^2^ / Conditional R^2^ | 0.062 / 0.646 | | 0.240 / 0.720 | | 0.295 / 0.718 | | 0.272 / 0.522 | | 0.276 / 0.673 | |

**Table S3. Summary of the linear mixed models between single trait community weighted means and intermittency, seasonal and connectivity factors.** The default modality of the DRN factor referred to the Croatian DRN. DOY refers to "day of the year’. ‘Connect.’ refers to spatio-temporal connectivity; “Drying dur.” to drying duration; “drying freq.” to drying frequency.

|  | **Aquatic passive dispersal** | | **Aquatic active dispersal** | | **Aerial passive dispersal** | | **Aerial active dispersal** | |
| --- | --- | --- | --- | --- | --- | --- | --- | --- |
| *Predictors* | *Coef* | *p* | *Coef* | *p* | *Coef* | *p* | *Coef* | *p* |
| (Intercept) | 2.05 | **<0.001** | 1.30 | **<0.001** | 0.55 | **<0.001** | 1.37 | **<0.001** |
| DRN [Czech] | -0.11 | **0.043** | -0.05 | 0.387 | 0.04 | 0.339 | 0.02 | 0.792 |
| DRN [Finland] | -0.24 | **<0.001** | -0.22 | **<0.001** | 0.19 | **<0.001** | 0.08 | 0.338 |
| DRN [France] | -0.04 | 0.451 | 0.02 | 0.658 | 0.00 | 0.953 | 0.10 | 0.252 |
| DRN [Hungary] | -0.34 | **<0.001** | -0.17 | **0.002** | -0.15 | **0.001** | -0.01 | 0.935 |
| DRN [Spain] | -0.10 | 0.082 | 0.10 | 0.071 | 0.19 | **<0.001** | 0.48 | **<0.001** |
| DOY | 0.00 | 0.985 | 0.02 | 0.918 | -0.39 | **0.013** | -0.50 | **0.018** |
| DOY:Drying dur. | -0.19 | **0.003** | -0.14 | **0.006** |  |  |  |  |
| DOY^2^ | -0.10 | 0.682 | 0.06 | 0.788 | -0.14 | 0.414 | 0.59 | **0.011** |
| DOY^2^:Drying dur. | -0.01 | 0.865 | -0.06 | 0.265 |  |  |  |  |
| Connect. |  |  | 0.06 | **0.005** | 0.06 | **0.013** | -0.00 | 0.968 |
| Connect.:Drying dur. |  |  |  |  |  |  | 0.04 | **0.002** |
| Connect.:Drying freq. |  |  |  |  |  |  | -0.06 | 0.146 |
| Drying dur. | -0.00 | 0.905 | 0.00 | 0.660 | 0.01 | 0.053 | -0.01 | 0.374 |
| Drying freq. | -0.09 | **<0.001** | -0.08 | **<0.001** |  |  | -0.04 | 0.293 |
| **Random Effects** | | | | | | | | |
| σ^2^ | 0.02 | | 0.01 | | 0.02 | | 0.03 | |
| τ00 | 0.02 site | | 0.02 site | | 0.01 site | | 0.06 site | |
| ICC | 0.49 | | 0.63 | | 0.41 | | 0.66 | |
| N | 126 site | | 126 site | | 126 site | | 126 site | |
| Observations | 636 | | 636 | | 636 | | 636 | |
| Marginal R^2^ / Conditional R^2^ | 0.359 / 0.671 | | 0.289 / 0.735 | | 0.365 / 0.626 | | 0.278 / 0.752 | |

**Table S4. Summary of the linear mixed models between single dispersal trait community weighted means and intermittency, seasonal and connectivity factors.** The default modality of the DRN factor referred to the Croatian DRN. DOY refers to ‘day of the year’. ‘Connect.’ refers to spatio-temporal connectivity; “Drying dur.” to drying duration; “drying freq.” to drying frequency.

|  | **Resistant eggs** | | **Cocoons** | | **Dormancy** | | **No resistance** | |
| --- | --- | --- | --- | --- | --- | --- | --- | --- |
| *Predictors* | *Coef* | *p* | *Coef* | *p* | *Coef* | *p* | *Coef* | *p* |
| (Intercept) | 0.51 | **<0.001** | 0.11 | **<0.001** | 0.42 | **<0.001** | 2.26 | **<0.001** |
| DRN [Czech] | 0.04 | 0.413 | 0.03 | 0.344 | 0.08 | **0.016** | -0.03 | 0.524 |
| DRN [Finland] | -0.14 | **0.005** | 0.12 | **<0.001** | 0.08 | **0.013** | -0.03 | 0.552 |
| DRN [France] | 0.10 | 0.057 | 0.10 | **0.004** | -0.04 | 0.246 | -0.00 | 0.950 |
| DRN [Hungary] | -0.06 | 0.271 | 0.17 | **<0.001** | 0.21 | **<0.001** | -0.28 | **<0.001** |
| DRN [Spain] | 0.25 | **<0.001** | -0.02 | 0.635 | -0.08 | **0.017** | -0.11 | **0.040** |
| DOY | -1.05 | **<0.001** |  |  | 0.05 | 0.804 | 0.35 | 0.230 |
| DOY:Drying dur. | -0.28 | **0.002** |  |  | 0.04 | 0.488 | -0.12 | 0.122 |
| DOY:Drying freq. | 1.09 | **<0.001** |  |  |  |  |  |  |
| DOY^2^ | 0.62 | **0.021** |  |  | 1.22 | **<0.001** | -1.08 | **0.001** |
| DOY^2^:Drying dur. | -0.05 | 0.559 |  |  | -0.14 | **0.010** | 0.37 | **<0.001** |
| DOY^2^:Drying freq. | -0.73 | **0.006** |  |  |  |  |  |  |
| Connect. | 0.01 | 0.681 | -0.00 | 0.898 |  |  | -0.06 | **0.032** |
| Connect.:Drying dur. | -0.02 | 0.069 |  |  |  |  |  |  |
| Connect.:Drying freq. | 0.15 | **<0.001** | -0.04 | **0.023** |  |  |  |  |
| Drying dur. | 0.02 | 0.113 | -0.02 | **0.006** | 0.02 | **<0.001** | -0.01 | **0.035** |
| Drying freq. | -0.11 | **<0.001** | 0.08 | **<0.001** |  |  |  |  |
| **Random Effects** | | | | | | | | |
| σ^2^ | 0.02 | | 0.01 | | 0.01 | | 0.03 | |
| τ00 | 0.02 site | | 0.01 site | | 0.01 site | | 0.02 site | |
| ICC | 0.53 | | 0.46 | | 0.33 | | 0.36 | |
| N | 126 site | | 126 site | | 126 site | | 126 site | |
| Observations | 636 | | 636 | | 636 | | 636 | |
| Marginal R^2^ / Conditional R^2^ | 0.345 / 0.689 | | 0.240 / 0.589 | | 0.384 / 0.587 | | 0.266 / 0.533 | |

### Table S5. Summary of the linear mixed models between single resistance trait community weighted means and intermittency, seasonal and connectivity factors. The default modality of the DRN factor referred to the Croatian DRN. DOY refers to ‘day of the year’. ‘Connect.’ refers to spatio-temporal connectivity; “Drying dur.” to drying duration; “drying freq.” to drying frequency.
